# Optogenetic modulation of TDP-43 oligomerization fast-forwards ALS-related pathologies in the spinal motor neurons

**DOI:** 10.1101/789057

**Authors:** Kazuhide Asakawa, Hiroshi Handa, Koichi Kawakami

## Abstract

Cytoplasmic aggregation of TDP-43 characterizes degenerating neurons in most cases of amyotrophic lateral sclerosis (ALS), yet the mechanisms and cellular outcomes of TDP-43 pathology remain largely elusive. Here, we develop an optogenetic TDP-43 variant (opTDP-43), whose multimerization status can be modulated *in vivo* through external light illumination. Using the translucent zebrafish neuromuscular system, we demonstrate that short-term light stimulation reversibly induces cytoplasmic opTDP-43 mislocalization, but not aggregation, in the spinal motor neuron, leading to an axon outgrowth defect associated with myofiber denervation. In contrast, opTDP-43 forms pathological aggregates in the cytoplasm after longer-term illumination and seeds non-optogenetic TDP-43 aggregation. Furthermore, we find that an ALS-linked mutation in the intrinsically disordered region (IDR) exacerbates the light-dependent opTDP-43 toxicity on locomotor behavior. Together, our results propose that IDR-mediated TDP-43 oligomerization triggers both acute and long-term pathologies of motor neurons, which may be relevant to the pathogenesis and progression of ALS.

## Introduction

Amyotrophic lateral sclerosis (ALS) is a neurological disorder in which the upper and lower motor neurons progressively degenerate, leading to muscular atrophy and eventually fatal paralysis. Trans-activation response element (TAR) DNA-binding protein 43 (TDP-43), a heterogeneous nuclear ribonucleoprotein, is mislocalized to the cytoplasm and forms pathological aggregates in the degenerating motor neurons in ALS ^1, 2^. TDP-43 aggregation characterizes almost all cases of sporadic ALS ^3, 4^, which accounts for greater than 90% of ALS. Moreover, mutations in the *TARDBP* gene encoding TDP-43 are linked to certain fraction (~ 4 %) of familial ALS ^5^. Despite its correlation with and causation of ALS, the role of TDP-43 in ALS pathogenesis has been largely unknown at the mechanistic level.

Multimerization of TDP-43 underlies its physiological and pathological roles. Under normal physiological conditions, homo-oligomerization of TDP-43 occurs through its N-terminal domain and is necessary for its RNA regulatory functions, such as splicing ^6–8^. At the C-terminus TDP-43 contains an intrinsically disordered region (IDR) with prion-like glutamine/asparagine-rich (Q/N) and glycine-rich regions, which can undergo liquid-liquid phase separation (LLPS) to form dynamic protein droplets ^9^. The TDP-43 IDR mutations that are linked to familial ALS cases enhance intrinsic aggregation propensity and protein stability of TDP-43 ^10 11^ and result in altered phase separation ^9^, which could contribute to disease propagation through acceleration of the formation and accumulation of pathological aggregates ^12, 13 14^. The modular architecture of TDP-43 has led to several hypotheses that its N-terminus-dependent oligomerization modulates C-terminal IDR-mediated aggregation either by enhancing ^9^ or hindering IDR interactions between adjacent TDP-43 molecules ^6 15^.

The severity of TDP-43 toxicity is correlated with the levels of wild-type and mutant TDP-43 expression in the various cellular and animal models ^16–25^. However, cytoplasmic TDP-43 aggregation is not always detectable in these models. Moreover, in a certain type of degenerating upper motor neurons, loss of nuclear TDP-43 can occur without the accumulation of cytoplasmic aggregates ^26^. Therefore, it has been difficult to evaluate how TDP-43 aggregation contributes to TDP-43 toxicity. Under these circumstances, it is necessary to develop a system to induce TDP-43 aggregation conditionally. Recently, light-dependent aggregation of *Arabidopsis* cryptochrome-2 was applied to the formation of IDR droplets via LLPS in a light illumination-dependent manner ^27^. This optogenetic approach has been successfully extended to the induction of cytotoxic TDP-43 aggregates formation in cultured cells ^28, 29^. However, interconversion of normal and toxic TDP-43 forms with spatiotemporal precision has not been achieved in animal models yet, which is central for the understanding of TDP-43 toxicity *in vivo*.

In the present study, we develop an optogenetic TDP-43 variant (opTDP-43) carrying a light-dependent oligomerization module of cryptochrome-2 attached to the IDR, and analyze the mechanisms of TDP-43 toxicity in spinal motor neurons *in vivo*. Transgenic expression and light stimulation of opTDP-43 in transparent zebrafish larvae show that oligomerization and aggregation of opTDP-43 is inducible and tunable by external light illumination *in vivo*. We reveal that, in the spinal motor neurons, short-term light illumination reversibly increases the cytoplasmic opTDP-43 pool and elevates myofiber denervation frequency in the absence of distinct aggregate formation. Furthermore, longer chronic light stimulation eventually leads to accumulation of cytoplasmic opTDP-43 aggregates that further seed aggregation of non-optogenetic TDP-43, which is accompanied by motor decline. The sequential pathological alterations of spinal motor neurons triggered by opTDP-43 oligomerization may provide clues about how motor neuron degeneration progresses at both molecular and cellular levels in a prodromal phase of ALS.

## Results

### Overexpression of TDP-43 causes cytoplasmic aggregation-independent toxicity in the spinal motor neurons

To explore mechanisms of TDP-43 toxicity associated with its cytoplasmic aggregation in spinal motor neurons, we first aimed to induce TDP-43 aggregation by its overexpression in the caudal primary motor neurons (CaPs) of zebrafish, which innervate a ventral third of the myotome and are present uniquely in every spinal hemisegment (Figure1A, B) ^30^. We generated a Gal4-inducible transgene of the zerbafish *tardbp*, encoding one of the two zebrafish TDP-43 paralogues, tagged with mRFP1 at its N-terminus (mRFP1-TDP-43z) (Figure 1C). To test the functionality of mRFP1-TDP-43z as TDP-43, we generated knock-out (KO) alleles for both *tardbp* and its paralogue *tardbpl* with the CRISPR-Cas9 system (i.e. *tardbp-n115* and *tardbpl-n94*, respectively). The TDP-43 double knock-out (DKO) embryos exhibited a blood circulation defect at 24-48 hours post-fertilization (hpf) (Sup. Figure 1D) and were lethal ^31^. We injected mRNA encoding wild-type Tardbp and mRFP1-TDP-43z into the TDP-43 DKO embryos at the one-cell stage (Sup. Figure 1A, B, C) and found that the mutant phenotype was rescued by both, indicating that mRFP1-TDP-43 is functional (Sup. Figure 1E). We then overexpressed mRFP1-TDP-43z in CaPs by combining Tg[UAS:mRFP1-TDP-43z] with the Tg[SAIG213A] driver (Figure 1A, B) ^32^, and analyzed their muscle innervation. The mRFP-TDP-43z overexpression significantly reduced the total axonal length at 48 hpf (Figure 1C, D, E, Sup Movie 1), while the axon arborized within the inherent innervation territory of the ventral myotomes (Figure 1C, D) and their branching frequency (i.e. branching as calculated per total axon length) were comparable to that of the wild-type CaP (Figure 1F), showing that overexpression of mRFP-TDP-43z primarily affects axon outgrowth, but not pathfinding or branching. However, the overexpressed mRFP-TDP-43z was predominantly accumulated in the nucleus and cytoplasmic aggregation was undetectable in the CaP at 48 hpf (Figure 1G). These observations suggest that an elevated level of TDP-43 causes neurotoxicity independently of cytoplasmic aggregation in the spinal motor neurons.

**Figure 1.**
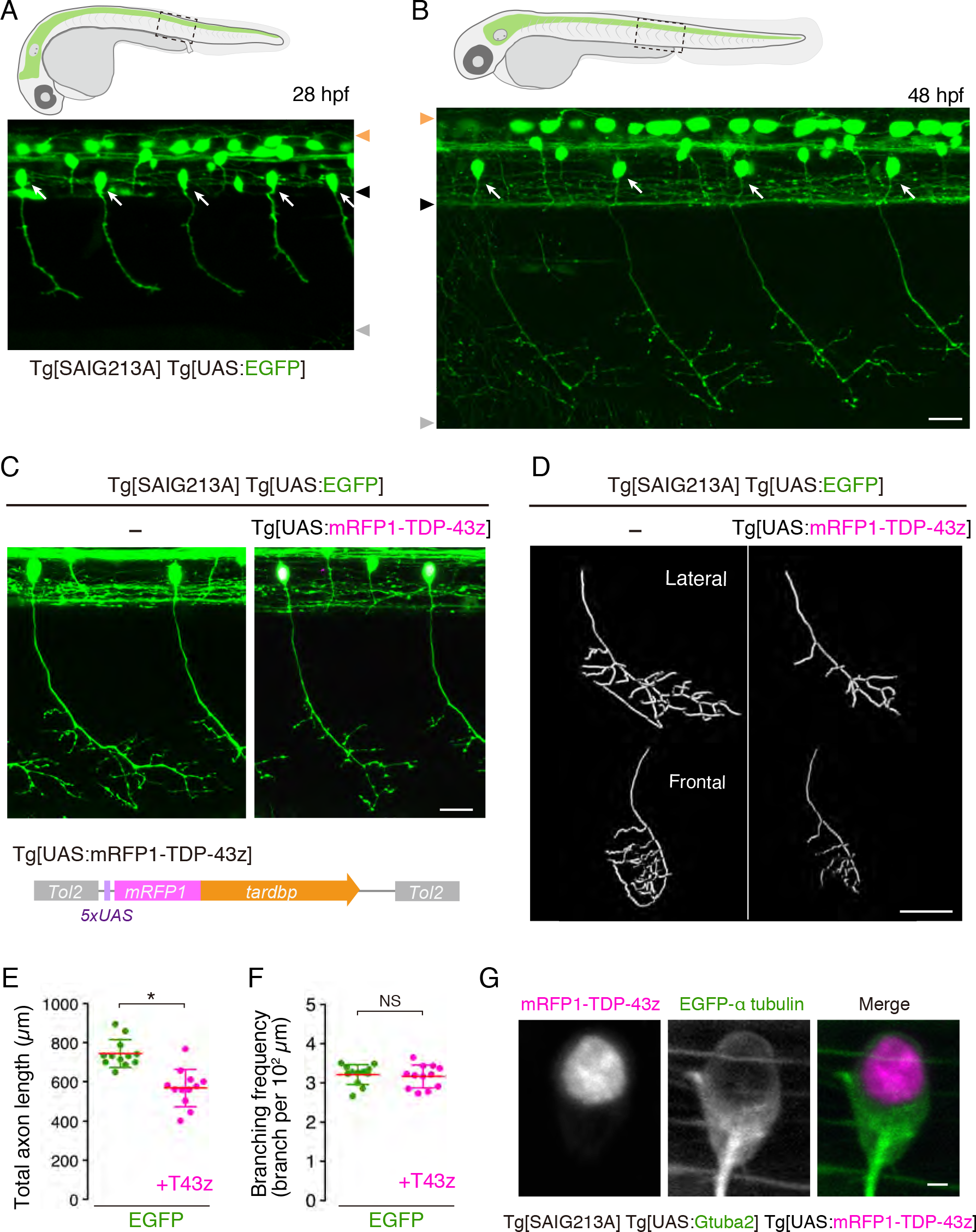
Overexpression of TDP-43 halts axon outgrowth independently of cytoplasmic TDP-43 aggregation. (A, B) CaPs (arrows) in Tg[SAIG213A] Tg[UAS:EGFP] fish. Orange, black and grey arrowheads indicate dorsal and ventral limits of the spinal cord, and ventral myotomal borders, respectively. (C) CaPs in Tg[SAIG213A] Tg[UAS:EGFP] (left) and Tg[SAIG213A] Tg[UAS:EGFP] Tg[UAS:mRFP1-TDP-43z] (right) larvae at 48 hpf. The structure of Tg[UAS:mRFP1-TDP-43z] transgene (bottom). (D) The lateral and frontal views of skeletonized CaP axons of Tg[SAIG213A] Tg[UAS:EGFP] (left) and Tg[SAIG213A] Tg[UAS:EGFP] Tg[UAS:mRFP1-TDP-43z] (right). The axon branch points and terminals are indicated by red and green, respectively. (See also Sup Movie1). (E, F) Total length and branching frequency of CaP axons at the spinal segment 14-17 of Tg[SAIG213A] Tg[UAS:EGFP] (green, 12 CaPs, 5 animals) and Tg[SAIG213A] Tg[UAS:EGFP] Tg[UAS:mRFP1-TDP-43z] (mageneta, 12 CaPs, 6 animals). *, p < 0.0001. NS, Not statistically significant. (G) Localization of mRFP1-TDP-43z in a CaP of Tg[SAIG213A] Tg[UAS:Gtuba2] Tg[UAS:mRFP1-TDP-43z] at 48 hpf. EGFP-taggedαtubulin expressed from Tg[UAS:Gtuba2] ^60^ serves as a marker for cytoplasm. The bars indicate 20 *μ*m (B, C), 40 *μ*m (D), 2 *μ*m (G).

### A photo-switchable TDP-43: opTDP-43

Next, we developed an alternative strategy to induce cytoplasmic TDP-43 aggregation. The IDR at the C-terminal of TDP-43 has a high propensity to form aggregates ^10^. Therefore, we reasoned that cytoplasmic TDP-43 aggregation might effectively occur when the proximity between IDRs was increased by addition of an exogenous multimerization tag, such as *Arabidopsis* cryptochrome CRY2. We first tested the feasibility of CRY2 oligomerization in spinal motor neurons *in vivo* via external light illumination. We created a transgenic zebrafish line carrying a fusion of mRFP1 and CRY2olig, a point mutant version of CRY2 (E490G) that exhibits significant clustering upon blue light illumination ^33^, under the UAS sequence (Figure 2A; Tg[UAS:mRFP1-CRY2olig]). In Tg[SAIG213A] Tg[UAS:mRFP1-CRY2olig] double transgenic embryos raised under dark conditions, mRFP1-CRY2olig was dispersed throughout the CaPs at 30 hpf (Figure 2B). Upon blue light illumination via confocal laser scanning of entire CaPs, mRFP1-CRY2olig instantaneously clustered in somas and axons during the first 10 min of illumination (Figure 2B, Sup Movie 2). Once illumination ceased, the mRFP1-CRY2olig clusters gradually disappeared and a homogeneous distribution of mRFP1-CRY2olig was restored (Figure 2B), showing that CRY2olig clustering is rapidly and reversibly controllable by light in the spinal motor neurons *in vivo*.

**Figure 2.**
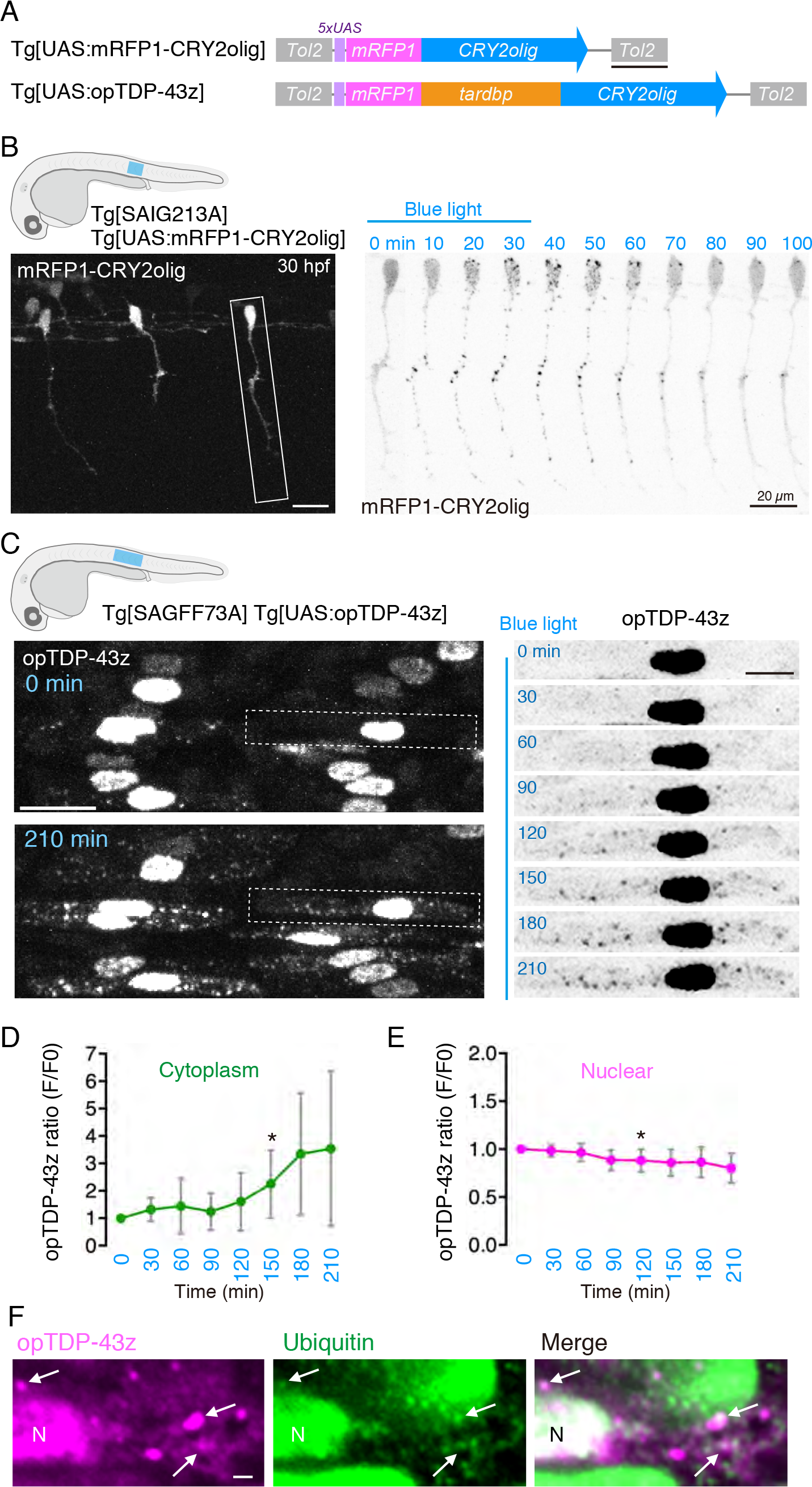
A photo-switchable TDP-43: opTDP-43. (A) The structures of Tg[UAS:mRFP1-CRY2olig] and Tg[UAS:opTDP-43z]. (B) Tg[SAIG213A] Tg[UAS:mRFP1-CRY2olig] fish at 30 hpf. A montage of a CaP expressing mRFP1-CRY2olig (boxed). The blue light was illuminated from 0 to 30 min. (C) Tg[SAGFF73A] Tg[UAS:opTDP-43z] fish at 28 hpf (0 min) and 31.5 hpf (210 hpf). A montage of a skeletal muscle cell expressing opTDP-43z (dashed box). The blue light was illuminated from 0 to 210 min. (D, E) The averaged change of opTDP-43 intensity in the cytoplasm (D) and nucleus (E) during the illumination (N = 8 cells). The asterisks indicate the emergence of statistically significant change in opTDP-43 fluorescence intensity in comparison to the opTDP-43 level at t = 0 (p < 0.05, t test). (F) Immunofluorescence of the skeletal muscle of the fish illuminated for 3.5 hours, using anti-RFP antibody (for opTDP-43) and anti-ubiquitin antibodies. Arrows indicate the representative of opTDP-43 droplets that are partially ubiquitinated. N, nucleus. The bars indicate 500 bp (A), 20 *μ*m (B, C left), 10 *μ*m (C right), and 2 *μ*m (F).

To adopt the clustering capacity of mRFP1-CRY2olig to TDP-43, we inserted the zebrafish *tardbp* between the mRFP1 and CRY2olig modules, and designated the resulting mRFP1-*tardbp*-CRY2olig fusion gene as opTDP-43z (i.e. optogenetic TDP-43 of zebrafish) (Figure 2A). The resulting opTDP-43z rescued the blood circulation defect of TDP-43DKO embryos under dark conditions as efficiently as wild-type *tardbp* (Sup Figure 1E), confirming that opTDP-43z is functional. We first assessed the igomerization capacity of optoTDP-43z in the skeletal muscle cells by taking advantage of their relatively large nucleus and cytoplasm. Since the strong whole body expression of mRFP-TDP-43z driven by the ubiquitous Gal4 driver Tg[SAGFF73A] perturbed development (Sup. Figure 2), we generated a UAS transgenic line that expressed a tolerable level of opTDP-43z with the Tg[SAGFF73A] driver (Figure 2A; Tg[UAS:opTDP-43z]). Unlike mRFP1-CRY2olig, opTDP-43z predominantly localized to the nucleus of the skeletal muscle cells under dark conditions (Figure 2C), suggesting that opTDP-43z localization is regulated by TDP-43-dependent mechanisms. We found that, while the nuclear-enriched opTDP-43z localization persisted during the 3.5 hours of blue light illumination (28-31.5 hpf), the cytoplasmic opTDP-43z gradually increased (Figure 2C, Sup Movie 3) and opTDP-43z droplets appeared 60-90 min after the initiation of illumination (Figure 2C, D). On the other hand, the nuclear opTDP-43z signal decreased slightly but significantly over time during the illumination (Figure 2E). We also found that the cytoplasmic opTDP-43z droplets were partially ubiquitinated as shown by immunofluorescence (Figure 2F), suggesting that the opTDP-43z level is regulated by proteolysis ^34, 35^. Altogether, these observations demonstrate that opTDP-43z is a photo-switchable variant of TDP-43 that forms aggregates in a blue light illumination-dependent manner.

### Light stimulation of opTDP-43z promotes cytoplasmic mislocalization in neuronal cells

To investigate light responsiveness of opTDP-43z in neuronal cells, we expressed opTDP-43z in both spinal motor neurons and tactile sensing Rohon-Beard (RB) cells by combining both Tg[mnr2b-hs:Gal4] ^36^ and Tg[SAIG213A] drivers. Under dark conditions, opTDP-43z primarily localized to the nucleus of both cell types at 28 hpf in Tg[mnr2b-hs:Gal4] Tg[SAIG213A] Tg[UAS:opTDP-43z] Tg[UAS:EGFP] quadruple transgenic fish (Figure 3A). Upon blue light illumination of the spinal cord, the nuclear-enriched localization of opTDP-43z persisted for about 90 min in both spinal motor neurons and RB cells, and then its localization was gradually expanded to the entire EGFP-positive area (Figure 3B C, Sup Movie 4), suggesting that light-dependent opTDP-43z oligomerization promotes its mislocalization to the cytoplasm. Unexpectedly, however, the cytoplasmic opTDP-43z mislocalization did not lead to distinct droplet formation as observed in the skeletal muscle cells within the time frames examined (up to 4.5 hours illumination), suggesting that the spinal motor neurons and RB cells have lower propensity to form opTDP-43 aggregates than the skeletal muscle cells. The cell type-dependent variation of opTDP-43 mislocalization and aggregation was also substantiated by the observations that neither embryonic epithelial cells nor differentiated skeletal muscle fibers displayed cytoplasmic mislocalization or aggregation of opTDP-43z under the same light illumination condition (Sup Figure 3).

**Figure 3.**
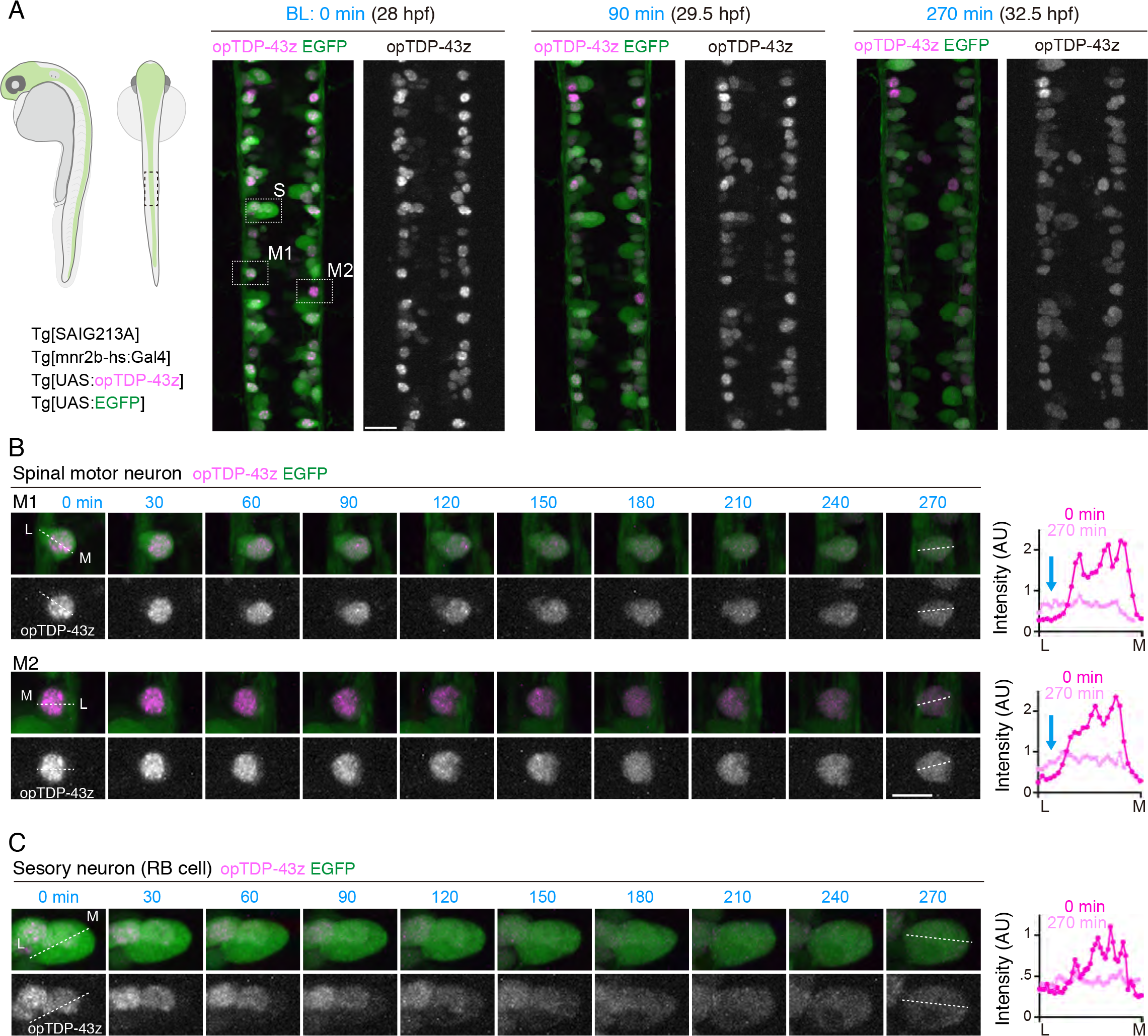
Light illumination-dependent cytoplasmic mislocalization of opTDP-43 in neuronal cells. **(A)** The dorsal view of the spinal cord at the segment 14-17 levels of a Tg[SAIG213A] Tg[mnr2b-hs:Gal4] Tg[UAS:opTDP-43z] Tg[UAS:EGFP] quadruple transgenic fish. Two spinal motor neurons (M1 and M2) and a Rohon-Beard sensory neuron (RB cell, S) highlighted with dashed boxes were analyzed in detail in B and C. (B, C) A montage of two spinal motor neurons (M1 and M2 in A) and a RB cell during the light illumination. The graphs show the fluorescent intensities of opTDP-43 along the dotted line drawn from the lateral (L) to medial edge (M) of the EGFP signal. The blue arrows indicate the cytoplasmic increase of opTDP-43. The bars indicate 20 *μ*m (A), 10 *μ*m (B).

### opTDP-43z oligomerization perturbs axon outgrowth

To explore the impact of light-induced opTDP-43z mislocalization at the whole cell level, we restricted opTDP-43z expression to CaPs by using the Tg[SAIG213A] driver. We devised a protocol by which Tg[SAIG213A] Tg[UAS:EGFP] Tg[UAS:opTDP-43z] triple transgenic fish were raised under continuous dark conditions until 48 hpf except being illuminated for 3 hours during 28-31 hpf (Figure 4A). Under this paradigm, opTDP-43z was primarily localized within the nucleus at 28 hpf, then dispersed throughout the nucleus and cytoplasm upon illumination, and restored its nuclear-enriched localization at 48-50 hpf (Figure 4A, B). We found by morphological analyses that total axon length, but not branching frequency, of CaPs decreased at 48-50 hpf by 13 % in the fish treated with 3 hours of blue light illumination, while such a phenotype was detected neither under continuous dark conditions nor by mRFP1-CRY2olig expression (Figure 4C, D, E). As observed with mRFP1-TDP-43z overexpression (Figure 1), the axons of light-stimulated CaPs expressing opTDP-43z arborized within their inherent ventral innervation territory (Figure 4C), and their branching frequency remained unchanged (Figure 4E), suggesting that the light-dependent opTDP-43z toxicity primarily influences axon outgrowth, but not pathfinding or branching.

**Figure 4.**
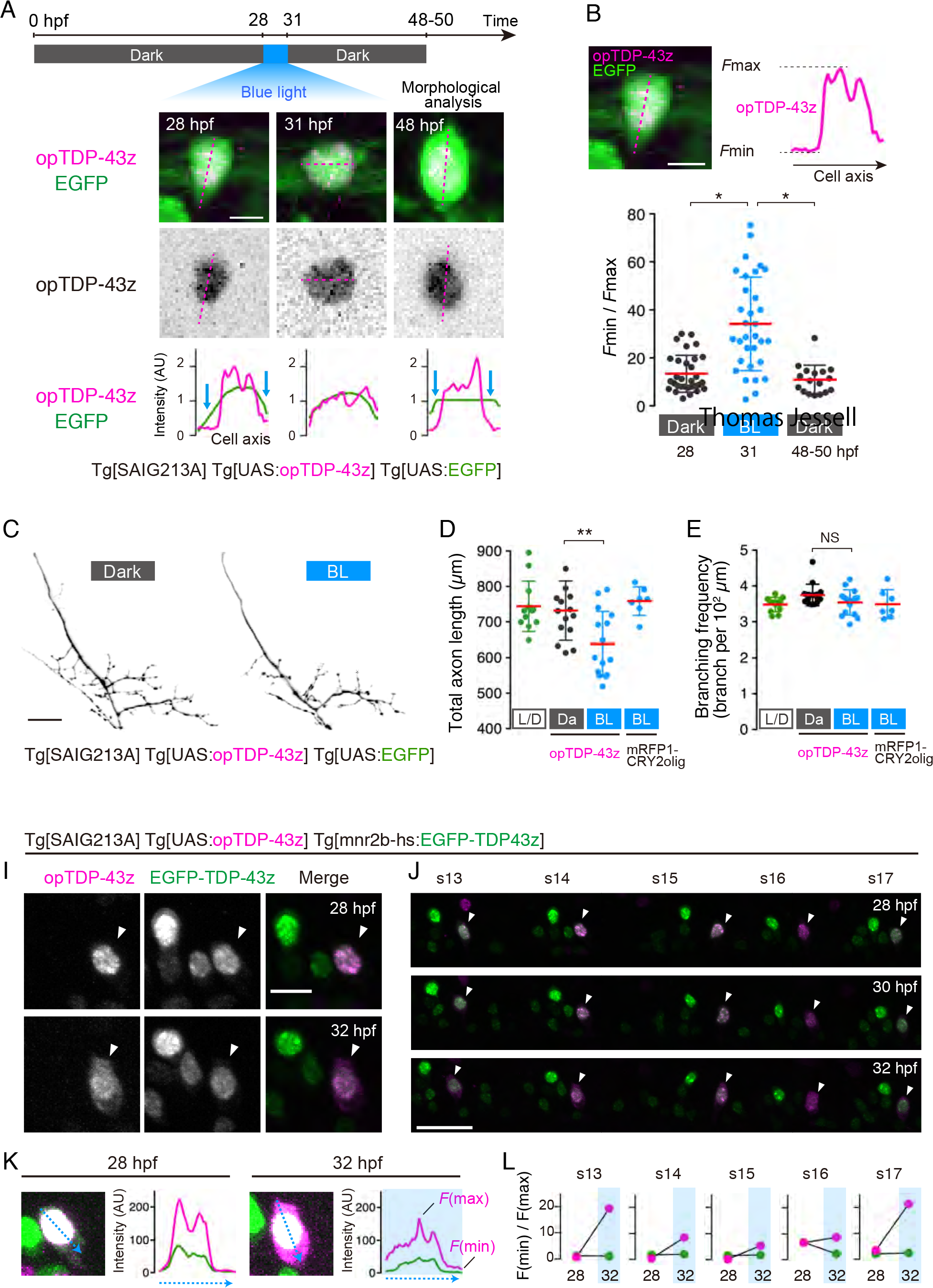
The light-dependent transient cytoplasmic mislocalization of opTDP-43z is accompanied by diminished motor axon outgrowth. (A) The light-illumination paradigm of CaPs. The spinal cord of Tg[SAIG213A] Tg[UAS:opTDP-43z] Tg[UAS:EGFP] fish at the spinal segment 13-18 level were illuminated with a blue laser, and CaPs were subjected to morphological analysis at 48-50 hpf. Shown images are a single CaP from dorsal (28 and 31 hpf) and lateral (48 hpf) views. The fluorescence intensity along the longest inner diameter (dashed magenta line) is plotted at each time point. Blue arrows indicate the presumptive cytoplasm area, where the opTDP-43z signal is faint. (B) Cytoplasmic shift of opTDP-43 localization is evaluated as a relative value of minimal (*F*min, cytoplasm) and maximal (*F*max, nuclear) fluorescence intensity along the longest inner diameter (dashed magenta line). The results were obtained from 32 cells (28 and 31 hpf) and 17 cells (48 hpf) in three independent Tg[SAIG213A] Tg[UAS:opTDP-43z] Tg[UAS:EGFP] fish. * p < 0.0001 (t test). (C) CaP motor axons with (BL) or without (Dark) blue light stimulation. (D, E) The total axon length and branching frequency of CaP motor axons in Tg[SAIG213A] Tg[UAS:EGFP] fish raised under normal laboratory light-dark cycle (L/D, the overlapping data sets with Figure 1E, F), Tg[SAIG213A] Tg[UAS:opTDP-43z] Tg[UAS:EGFP] fish with (BL, 15 cells, 4 animals) or without (Da, 15 cells, 4 animals) blue light stimulation, and Tg[SAIG213A] Tg[UAS:mRFP1-CRY2olig] Tg[UAS:EGFP] fish with the blue light stimulation (7 cells, 2 animals). ** p = 0.0068 (t test). NS, not significant. (I, J) Somas of the CaPs (arrowhead) and other *mnr2b*-positive spinal motor neurons in the spinal segment 14 (I) and 13-17 (J) of Tg[SAIG213A] Tg[UAS:opTDP-43z] Tg[mnr2b-hs:EGFP-TDP43z] fish that was illuminated with a blue light during 28 hpf-32 hpf. (K, L) Evaluation of cytoplasmic shift of opTDP-43z and EGFP-TDP-43z in the CaP in I. (K) The fluorescence intensities of opTDP-43z (magenta) and EGFP-TDP-43z (green) was plotted along the blue dashed arrows. Images shown are over enhanced for identification of soma outline. (L) The relative intensity of cytoplasmic signal (*F*min/*F*max) for opTDP-43z (magenta) and for EGFP-TDP-43z (green) in each spinal segment. The bars indicate 5 *μ*m (A, B), 20 *μ*m (C, J), 10 *μ*m (I).

The absence of distinct cytoplasmic aggregate formation in the CaPs raises the possibility that opTDP-43z exerts its toxicity through dragging of non-optogenetic TDP-43 out of the nucleus to the cytoplasm. To test this possibility, we constructed Tg[mnr2b-hs:EGFP-TDP-43z] expressing EGFP-TDP-43z under the control of the mnr2b promoter, and established a Tg[SAIGFF213A] Tg[UAS:opTDP-43z] Tg[mnr2b-hs:EGFP-TDP-43z] triple transgenic fish. Under dark conditions at 28 hpf, both opTDP-43z and EGFP-TDP-43z exhibited nuclear localization with subnuclear distribution patterns similar to each other (Figure 4I). Contrary to our prediction, however, the nuclear-enriched EGFP-TDP-43z localization remained unaffected while opTDP-43z was dispersed throughout the soma with 4 hour-blue light illumination during 28-32 hpf (Figure 4I-K), demonstrating that light-induced opTDP-43z mislocalization occurs independently of the non-optogenetic TDP-43 pool. These observations suggest that the perturbation of axon outgrowth by light-stimulated opTDP-43z is unlikely to be caused by loss of TDP-43 function due to nuclear TDP-43 reduction or depletion.

### Light-stimulated opTDP-43z elevates myofiber denervation frequency

The opTDP-43z-mediated axon outgrowth defects raised the question as to whether opTDP-43z perturbs axon extension or promotes axon shrinkage, or both. To address this, we analyzed a major axon collateral of CaP innervating the dorsal side of its innervation territory that had experienced tertial branching at 56 hpf (provisionally named *d*orsal axon *c*ollateral of *C*aP with *t*ertial branching: DCCT) (Figure 5A). Live imaging revealed that the total DCCT length increased by 26 % from 56 to 72 hpf, (Figure 5B, C) and formed one additional branch on average in Tg[SAIG213A] Tg[UAS:GFP] larvae. Intriguingly, we noticed that a minor but significant population of single CaPs (24 %) increased their total DCCT length with a decrease in the number of collateral branches (Figure 5D), indicating that normal DCCT outgrowth involves both extension and shrinkage, as the extension occurs more frequently. The expression of opTDP-43z itself did not affect the average DCCT outgrowth rate and branch number under dark conditions at 56 hpf (Figure 5D, E). On the other hand, the average DCCT growth rate significantly declined by 11 % with at 72 hpf, when CaPs expressing opTDP-43z had been illuminated for 3 hours (from 56 to 59 hpf) (Figure 5D). Remarkably, 28 % (5 out of 18 DCCTs) of the illuminated CaPs exhibited total DCCT lengths (Figure 5C), and 44 % showed reduced DCCT branch number (Figure 5E), demonstrating that axon shrinkage contributes to the observed axon outgrowth defects.

**Figure 5.**
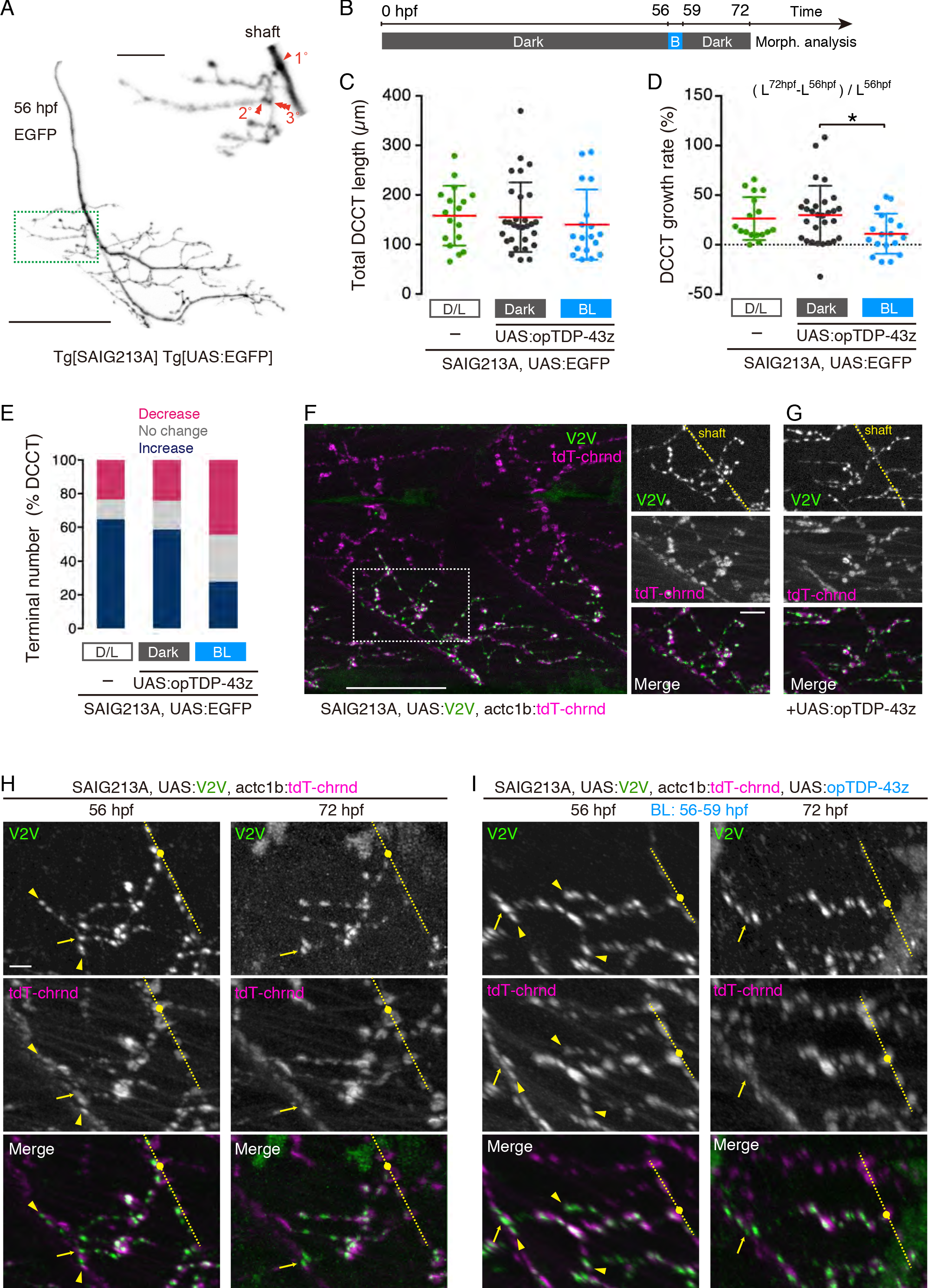
Axonal shrinkage and myofiber denervation caused by light stimulation of opTDP-43z. (A) A CaP motor axon in Tg[SAIG213A] Tg[UAS:EGFP] fish. DCCT (green box) was magnified on the right. Primary, secondary and tertial branchings were indicated in red. (B) Light-illumination paradigm. (C, D, E) Total length (C), growth rate (D) and population change in axon terminal number (E) of DCCTs in Tg[SAIG213A] Tg[UAS:EGFP] Tg[UAS:opTDP-43z] fish (Dark: 29 cells, 5 animals, BL: 18 cells, 3 animals) and Tg[SAIG213A] Tg[UAS:EGFP] fish raised under normal dark light cycles (D/L: 17 cells, 3 animals). *, p = 0.0224 (t test). (F) The lateral view of the trunk of Tg[SAIG213A] Tg[UAS:V2V] Tg[actc1b:tdT-chrnd] fish (left) and neuromuscular synapses of the DCCT (right) at 56 hpf. The dashed yellow lines indicate the CaP axon shaft. (G) Neuromuscular synapses of the DCCT in Tg[SAIG213A] Tg[UAS:opTDP-43z] Tg[UAS:V2V] Tg[actc1b:tdT-chrnd] fish at 56 hpf. (H, I) Live imaging of DCCT neuromuscular synapses. Yellow arrowheads indicate the neuromuscular synapses that were not present at 72 hpf. The yellow dashed lines, dots, arrows indicate axon shafts, primary branching points, and contact sites with the myotomal boundaries of CaPs, respectively. In I, Z-stacks are produced from 3D-rotated images made by Imaris, to make the denervation events (arrowhead) clearly visible. The bars indicate 10 *μ*m (A top, F right), 50 *μ*m (A bottom, F left), 5*μ*m (H).

We then investigated whether the DCCT shrinkage involves myofiber denervation by live monitoring of pre- and postsynaptic structures with Vamp2-Venus and tdTomato-tagged acetylcoline receptor (dT-chrnd), respectively (Figure 5F) ^36^. We found that, in both wild-type and opTDP-43z expression conditions prior to light stimulation, the DCCT axon terminals were decorated by Vamp2-Venus and the Vamp2-Venus signals were well colocalized with dT-chrnd (Figure 5F, G), indicating normal neuromuscular assembly (Figure 5F). Live imaging revealed that, in the opTDP-43z-expressing CaPs after the 3 hours of illumination (during 56-59 hpf), Vamp2-Venus and juxtaposed dT-chrnd speckles at the axon terminal disappeared (Figure 5I), demonstrating that a decrement in DCCT terminal number is accompanied by myofiber denervation (Figure 5E). Such disappearance of juxtaposed Vamp2-Venus and dT-chrnd was also observed in wild-type DCCTs (Figure 5E, H). Overall, these observations show that the DCCT shrinkage is associated with myofiber denervation, and that optogenetic TDP-43 oligomerization raises the denervation frequency.

### Cytoplasmic aggregation of opTDP-43h that seeds non-optogenetic TDP-43 aggregation in the spinal motor neurons

Targeted optogenetic stimulation via confocal laser scanning required the fish to be agarose-embedded, which restricted the illumination duration to a maximum of ~4 hours fully maintain fish viability. We aimed to test if a longer illumination period potentially induces cytoplasmic aggregation of the optogenetic TDP-43 in the spinal motor neurons. We constructed transgenic fish in which most of the spinal motor neurons expressed a CRY2olig-tagged human TDP-43 (opTDP-43h, for optogenetic TDP-43 of human) from an mnr2b-BAC transgene (Tg[mnr2b-hs:opTDP-43h]) (Figure 6A)(Sup. Figure 1E)^31^ and established a system for longitudinal field illumination of blue LED light using unrestrained fish. The Tg[mnr2b-hs:opTDP-43h] and Tg[mnr2b-hs:EGFP-TDP-43z] transgenes were combined to allow for simultaneous live monitoring of opTDP-43h and non-optogenetic TDP-43 in the spinal motor neurons. Prior to the illumination, both opTDP-43h and EGFP-TDP-43z were primarily localized in the nucleus at 2 days post-fertilization (dpf) (Figure 6B). We found that opTDP-43h was dispersed throughout the cell and formed aggregates in the cytoplasm at 3 dpf, and that aggregation was further enhanced over the subsequent 48 hours of illumination (i.e. 3-5 dpf). Despite distinct cytoplasmic opTDP-43h mislocalization and aggregation, EGFP-TDP-43z was predominantly localized to the nucleus during 2-4 dpf, suggesting that opTDP-43h mislocalization and aggregation occurred independently of EGFP-TDP-43z during the 48 hours of illumination. Intriguingly, at later time points (e.g. 5 dpf) when the cytoplasmic opTDP-43h developed into larger aggregates, EGFP-TDP-43z became detectable in the cytoplasm and formed distinct foci that colocalized with opTDP-43h aggregates (Figure 6C), indicating that, the long term light-induced opTDP-43h aggregates seeds aggregation of non-optogenetic TDP-43.

**Figure 6.**
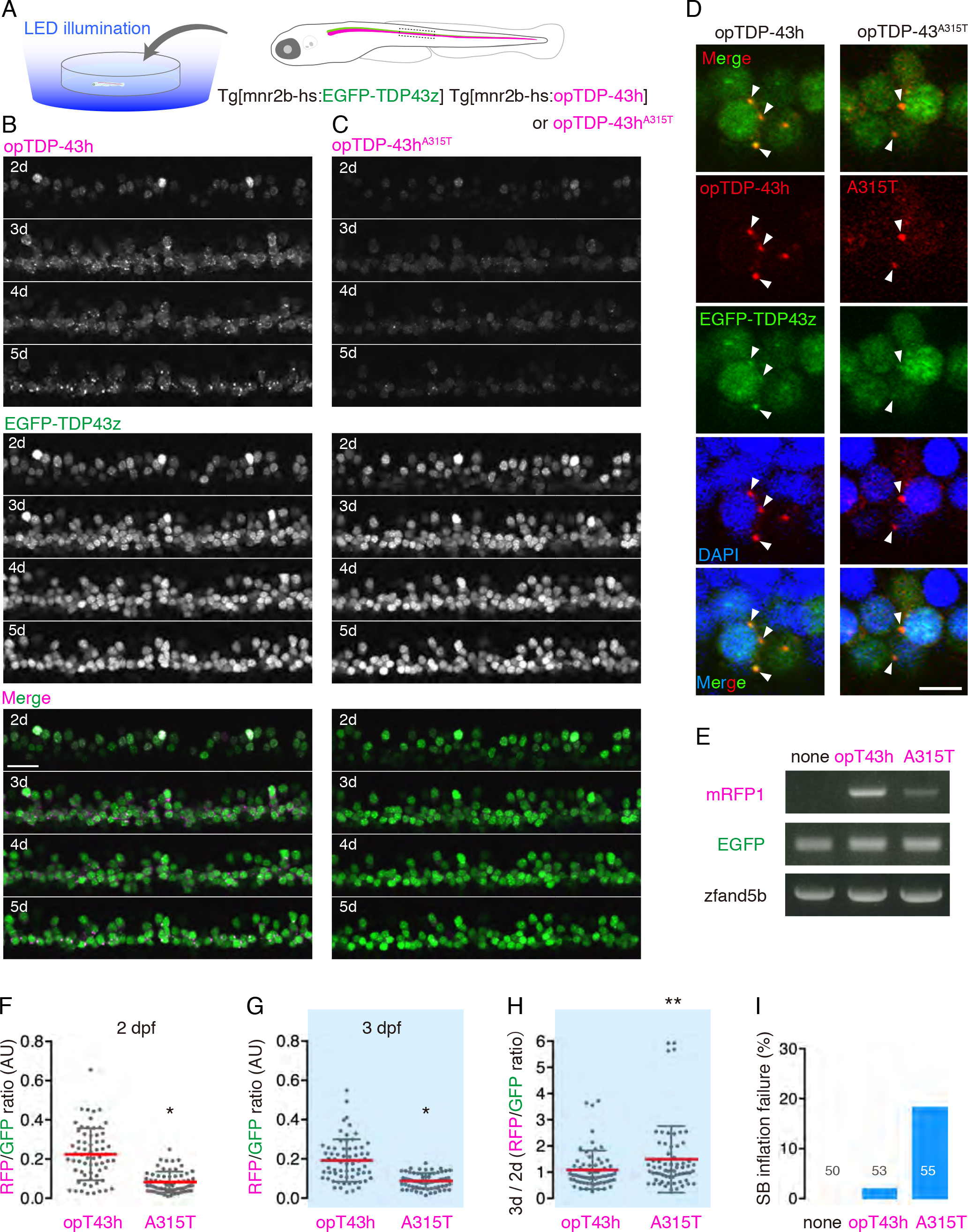
Long-term light stimulation of opTDP-43h induces TDP-43 aggregation that seeds non-optogenetic TDP-43 aggregation. (A) Chronic field illumination of unrestrained larvae from 2 dpf -5 dpf. (B, C) Live imaging of the spinal motor column of Tg[mnr2b-hs:EGFP-TDP43z] Tg[mnr2b-hs:opTDP-43h] (B) and Tg[mnr2b-hs:EGFP-TDP43z] Tg[mnr2b-hs:opTDP-43h^A315T^] (C) fish from 2 - 5 dpf. (D) Chronically light-stimulated opTDP-43h (left) and opTDP-43h^A315T^ (right) aggregates in the cytoplasm and seed EGFP-TDP-43z aggregation. Arrowheads indicate opTDP-43 and opTDP-43h^A315T^ aggregates that contain EGFP-TDP-43z. (E) RT-PCR analysis for opTDP-43h (opT43h) and opTDP-43h^A315T^ (A315T). cDNA samples were extracted from Tg[mnr2b-hs:EGFP-TDP43z] (none), Tg[mnr2b-hs:EGFP-TDP43z] Tg[mnr2b-hs:opTDP-43h] (opT43h), and Tg[mnr2b-hs:EGFP-TDP43z] Tg[mnr2b-hs:opTDP-43h^A315T^] (A315T) larvae at 72hpf. EGFP and zfand5b are internal controls. (F, G, H) Longitudinal single cell analyses of opTDP-43h or opTDP-43h^A315T^ fluorescence intensity (RFP) relative to EGFP-TDP-43z (GFP) (64 cells from 4 animals, for each). * p < 0.001 (t test). ** p = 0.03 (t test). (I) Failure rate of swimming bladder (SB) inflation of Tg[mnr2b-hs:EGFP-TDP43z] (none), Tg[mnr2b-hs:EGFP-TDP43z] Tg[mnr2b-hs:opTDP-43h] (opT43h), and Tg[mnr2b-hs:EGFP-TDP43z] Tg[mnr2b-hs:opTDP-43h^A315T^] (A315T) larvae at 5 dpf. Figures indicate the number of illuminated fish examined. SB inflation failure was not observed when fish were raised under normal dark light cycles (N > 100 for each). The bars indicates 20 *μ*m (B) and 5 *μ*m (D).

### IDR mutation A315T enhances protein stability and oligomerization-dependent toxicity of TDP-43

The gradual mislocalization and aggregation of non-optogenetic TDP-43 promoted by the light-stimulated opTDP-43h prompted us to hypothesize that opTDP-43h first oligomerizes via the CRY2olig module and subsequently seeds non-optogenetic TDP-43 aggregation, via its aggregate-prone IDR ^10, 37^. To test whether the IDR contributes to light-dependent toxicity of opTDP-43h, we created an opTDP-43 mutant with an IDR mutation (A315T) linked to familial ALS (opTDP-43h^A315T^) and expressed opTDP-43h^A315T^ widely in the spinal motor neurons from Tg[mnr2b-hs:opTDP-43h^A315T^]. In Tg[mnr2b-hs:opTDP-43h^A315T^] Tg[mnr2b-hs:EGFP-TDP-43z] double transgenic fish, opTDP-43h^A315T^ displayed nuclear-enriched localization prior to the LED illumination at 48 hpf. We noted that the expression level of opTDP-43h^A315T^ protein was less than that of opTDP-43h in the Tg[mnr2b-hs:opTDP-43h] fish, partly due to the lower level of mRNA (Figure 6E). Nonetheless, in response to illumination, opTDP-43h^A315T^ mislocalized to the cytoplasm, formed aggregates within 24 hours of illumination and seeded cytoplasmic aggregates containing non-optogenetic EGFP-TDP-43z (Figure 6C, D). We then quantified the amounts of opTDP-43h and opTDP-43h^A315T^ proteins in the single cells before and after the initial 24 hours of illumination (2-3 dpf), by using EGFP-TDP-43z expressed from Tg[mnr2b-hs:EGFP-TDP-43z] as an internal control. the relative intensity of opTDP-43h^A315T^ significantly increased during the illumination by 3 dpf while that of opTDP-43h remained unchanged (Figure 6F, G, H), indicating that the A315T mutation increases the protein stability. This observation is consistent with the previous observations in cultured neurons ^11, 38^. The illuminated Tg[mnr2b-hs:opTDP-43h] Tg[mnr2b-hs:EGFP-TDP-43z] larvae were viable with seemingly normal free-swimming activity at 5-6 dpf, suggesting that the toxicity associated with light-stimulated opTDP-43h has only a minor effect at the behavioral level. On the other hand, 18 % of the illuminated fish expressing opTDP-43h^A315T^, but none of the non-illuminated siblings, failed to inflate the swim bladder at 5 dpf and showed declined locomotor ability (Figure 6I, Sup Movie 5), indicating that the A315T mutation enhances oligomerization-dependent toxicity of opTDP-43h. Altogether, these observations suggest that the IDR of TDP-43 causes the oligomerization-dependent toxicity in the spinal motor neurons.

## Discussion

Pathological aggregation of TDP-43 via the IDRs is proposed to be antagonized by N-terminal-mediated homo-oligomerization under physiological conditions ^6, 15^. In this study, we successfully developed CRY2olig-mediated TDP-43 oligomerization system *in vivo* and demonstrated CRY2olig-mediated oligomerization led to the accumulation of cytoplasmic opTDP-43 aggregates in the zebrafish spinal motor neurons. This CRY2olig-driven opTDP-43 oligomerization would initially generate reversible interactions within the IDRs, some of which occasionally transform into an irreversible form. Then, such irreversible “knots” of opTDP-43 eventually seed IDR-mediated aggregation of non-optogenetic TDP-43 (Figure 7). Under our illumination conditions, the spinal motor neurons require up to 3 hours to fully disperse opTDP-43 throughout the cell, 24 hours to accumulate distinct cytoplasmic opTDP-43 aggregates, and several additional days to develop cytoplasmic opTDP-43 aggregates containing non-optogenetic TDP-43. This sequentially regulated illumination-triggered TDP-43 knot and aggregate formation enables direct observation of spinal motor neuron pathology as triggered by IDR-mediated TDP-43 oligomerization. We propose that this opTDP-43-triggered pathology may correspond to a fast-forwarding of spinal motor neuron degeneration in ALS, in which a majority of cases are believed to involve IDR-mediated TDP-43 aggregation, yet currently allows very restricted anatomical access.

**Figure 7.**
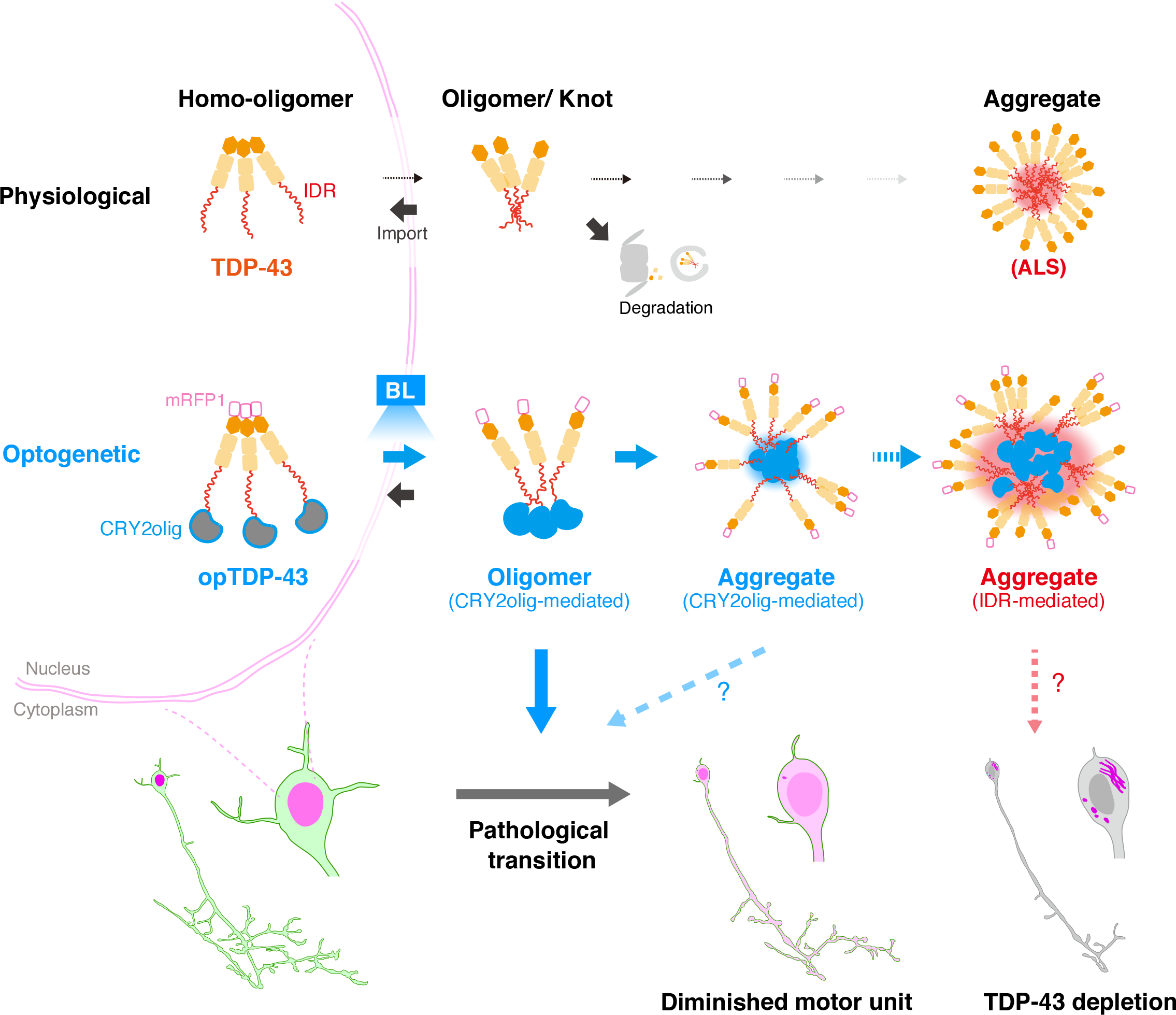
Sequentially regulated illumination-triggered TDP-43 Knot and Aggregate formation. In physiological conditions, TDP-43 forms oligomers via its N-terminus and is primarily localized in the nucleus. Spinal motor neurons keep cytoplasmic concentration of TDP-43 oligomers at a low level to prevent them from turning into toxic irreversible oligomers mediated by the C-terminus IDRs (toxic “knots”), which possess competence for developing into pathological TDP-43 aggregates, a hallmark of ALS. CRY2olig-driven opTDP-43 oligomerization promotes pathological change of the motor neurons, such as axon retraction associated with myofiber denervation, prior to accumulation of distinct cytoplasmic aggregates. Whether CRY2olig-diriven opTDP-43 aggregates are toxic to the motor neurons and whether CRY2olig-diriven aggregates eventually deplete endogenous nuclear TDP-43 pools are unknown.

We discovered that the reversible cytoplasmic opTDP-43 mislocalization induced by short-term light illumination was sufficient to cause defective motor axon outgrowth accompanied by enhanced myofiber denervation. The physiological nuclear-enriched TDP-43 localization is sustained by nucleocytoplasmic transport system ^34, 35^ as well as protein degradation systems in the cytoplasm, such as the ubiquitin proteasome system (UPS) and autophagy ^39, 40^. Since the light-dependent cytoplasmic opTDP-43 mislocalization commences without affecting non-optogenetic TDP-43 localization, the toxicity accompanied by opTDP-43 mislocalization may not be caused by a global shutdown of nucleocytoplasmic transport or proteolysis systems for TDP-43, but rather by dysregulation of RNAs and/or proteins being bound by opTDP-43. TDP-43 can associate with more than 6,000 RNA targets ^41–44^, and RNA-binding is antagonistic to toxic TDP-43 oligomerization in an optogenetic cellular model ^28^, implying that light-dependent opTDP-43 oligomerization would profoundly affect its RNA-binding capacity, thereby influencing the expression of a myriad of genes. In terms of the axon outgrowth defect, whether toxicity is ascribable to dysregulation of specific key proteins ^39, 45^ or to a widespread translational abnormality ^46^, which could lead to stress-inducing misfolded protein accumulation, remains to be investigated. Nevertheless, the normal motor axon pathfinding and unaffected branching frequency suggest a certain specificity of opTDP-43 toxicity and would favor the idea that the cellular growth pathway is primarily affected by the toxicity. It should also be noted that our current illumination paradigm encompasses not only the somas but also the motor axons. Therefore, “resident” cytoplasmic opTDP-43, such as that included in mRNP granules undergoing axonal transport ^47^ and in mitochondria at the axon terminals ^48, 49^, could also be photoconverted *in situ*, thereby contributing to the acute toxicity that involves neuromuscular synapse destabilization. Spatially-resolved light stimulation, a major advantage of optogenetics, could identify such potentially multiple pathogenic origins in the future.

The light-dependent cytoplasmic opTDP-43 mislocalization provides unexpected insight into the relationships between TDP-43 multimerization, localization, and toxicity, given that a toxic level of mRFP1-TDP-43 overexpression led neither to cytoplasmic mislocalization nor aggregation. The persistent nuclear localization of overexpressed mRFP1-TDP-43 indicates a robustness of the nucleocytoplasmic transport and cytoplasmic protein degradation systems against proteostatic perturbation of TDP-43. On the other hand, these TDP-43 surveillance systems appear to be inert against the IDR-mediated TDP-43 oligomers, as evidenced by the light-dependent cytoplasmic opTDP-43 mislocalization. These manipulations of proteostasis and multimerization revealed that a dosage increase of TDP-43 does not immediately lead to IDR-mediated oligomerization in the spinal motor neurons, and therefore TDP-43 toxicity associated with proteostatic abrogation could be mechanistically distinct from that caused by IDR-mediated oligomerization. Importantly, our animal model approach explicitly revealed a striking cell-type variation of opTDP-43 mislocalization and aggregation, such that the neuronal cells are less prone to accumulate opTDP-43 aggregates compared to the differentiating muscle cells, while both of these cell types are inherently more competent for cytoplasmic mislocalization than the epithelial cells. Although the mechanisms underlying this cell-type specificity remain unknown, the present and previous observations ^50^ emphasize the importance of studying vulnerable cell types *in vivo* for accurately disclosing the mechanisms underlying TDP-43 localization, and thereby toxicity.

The verification that CRY2olig-mediated opTDP-43 oligomerization is toxic to the spinal motor neurons instead made it difficult to evaluate the toxicity derived from opTDP-43 aggregates alone, as the oligomeric and aggregate forms of opTDP-43 coexists during illumination. In this regard, it is noteworthy that the opTDP-43h^A315T^ that was expressed less and formed fewer cytoplasmic aggregates was more toxic than the opTDP-43h expressed in larger amounts (Figure 6), which provides an *in vivo* example in which the amount of accumulated TDP-43 aggregates does not necessarily predict the degree of TDP-43 toxicity ^16, 17^. It was recently proposed that TDP-43 adopts both reversible and irreversible β-sheet aggregates that are involved in the formation of membraneless organelles, such as stress granules (SGs) and pathogenic amyloids, respectively, and that ALS mutations, including A315T, can promote the transition of such reversible to irreversible pathogenic aggregation ^13^. Also, in frontotemporal lobar degeneration (FTLD), TDP-43 displays distinct aggregate assemblies and toxic effects in disease-subtype-specific manners ^51^. Thus, it is possible that light-induced opTDP-43 aggregates are in fact heterogeneous and only a certain form of aggregates become pathogenic and acquire seeding activity for non-optogenetic TDP-43 aggregation. The time lag (~48 hours) between opTDP-43’s own aggregation and subsequent opTDP-43-dependent non-optogenetic TDP-43 aggregation might represent the time required for speciation of IDR interactions to develop such seeding capacity. The toxicity of opTDP-43 aggregates should be evaluated by considering such potential heterogeneity, and therefore remains as a challenging but important question to be addressed. Alternatively, the seeding capacity for non-optogenetic TDP-43 aggregation suggests that the toxicity of opTDP-43 aggregates can be exerted as a long-term effect through gradual depletion of the available nuclear and cytoplasmic TDP-43 pools into these aggregates, which would manifest as a TDP-43 loss-of-function phenotype.

It has been estimated that, even during healthy aging, the spinal motor neurons are substantially lost ^52–54^. As a result, the surviving motor units are enlarged to preserve maximal force generating capacity by compensatory collateral reinnervation ^55^. ALS has also been characterized by an elevated number of muscle fibers innervated by a single subterminal axon ^56^, which is likely a remnant of such compensatory collateral reinnervation events. In the present study, we found by live imaging of axon collateral that an innervation territory of healthy spinal motor neurons is determined by a balance between assembly and disassembly of neuromuscular synapses in zebrafish. We further discovered that optogenetic opTDP-43 oligomerization could tip the balance toward disassembly and decrease the total collateral length. Therefore, our results predict that, once a cellular concentration of IDR-mediated TDP-43 oligomers reaches a critical level, a spinal motor neuron would begin to reduce its motor unit size through repetition of incomplete denervation/reinnervation cycles. Such neurons would also be defective in complementing damaged neighboring motor units through collateral reinnervation, which would accelerate the manifestation of motor decline. We envision that opTDP-43 allows for approaching the mechanisms underlying such dynamic innervation/reinnervation balancing of spinal motor axons in health and TDP-43-associated pathology, as well as for interrogating how not only motor neurons but also diverse types of surrounding cells, including muscle, glial and endothelial cells, respond to and modify TDP-43 toxicity. Moreover, combined with the feasibility of high-throughput, whole organism chemical screening in zebrafish, opTDP-43-mediated motor neuron pathogenesis should be extended for exploring small molecules that restore a normal denervation/reinnervation balance for spinal motor neurons, which might serve as drugs for ALS and other TDP-43 proteinopathy.

## Acknowledgements

Authors would like to thank Drs Keiko Imamura and Haruhisa Inoue for valuable discussions and Kawakami lab members for generous support. This work was supported by SERIKA FUND (KA), The Nakabayashi Trust For ALS Research (KA), THE KATO MEMORIAL TRUST FOR NAMBYO RESEARCH (KA), Daiichi-Sankyo Foundation of Life Science (KA), Takeda Science Foundation (KA), KAKENHI Grant numbers JP16K07045 (KA), JP19K06933 (KA), National BioResource Project from Japan Agency for Medical Research and Development (AMED) (K.K.), and KAKENHI Grant Numbers JP15H02370 (K.K.).

## Author contributions

KA conceived the research, designed and performed the experiments. KA and KK analyzed the data and wrote the manuscript with inputs from HH.

## Methods

### Fish lines

This study was carried out in accordance with the Guide for the Care and Use of Laboratory Animals of the Institutional Animal Care and Use Committee (IACUC, approval identification number 24-2) of the National Institute of Genetics (NIG, Japan), which has an Animal Welfare Assurance on file (assurance number A5561-01) at the Office of Laboratory Animal Welfare of the National Institutes of Health (NIH, USA). Fish were raised under 12:12 light-dark (L/D) cycles during the first 5 days after birth, unless otherwise stated.

#### Transgenic fish lines

Tg[UAS:mRFP1-TDP-43z] was generated by synthesizing a *Tol2* transposon-based cassette (UAS:mRFP1-TDP-43z) carrying the zebrafish *tardbp* (Genbank accession # NM_2014476) that was tagged with mRFP1 (Genbank accession # AF506027.1) at the N-terminus with linker peptide and placed downstream of x5 upstream activation sequence (UAS)^57^ (pBMH-T2ZUASRzT43, Biomatik). For constructing Tg[UAS:mRFP1-CRY2olig], the zebrafish codon-optimized photolyase homology region (PHR) of *Arabidopsis thaliana* CRY2 carrying the E490G mutation (CRY2olig)^33^ was synthesized (Biomtik) and N-terminally tagged with mRFP1 with a linker peptide TRDISIE encoded by ACG CGT GAT ATC TCG ATC GAG (mRFP1-CRY2olig). The mRFP1-CRY2olig fragment was fused to 5xUAS, cloned into the Tol2-transposon cassette. For the construction of Tg[UAS:opTDP-43z], the mRFP1-TDP-43z fragment was C-terminally fused to CRY2olig with the linker peptide TRDISIE (opTDP-43z). The opTDP-43z fragment was fused to 5xUAS, cloned into the Tol2-transposon cassette. NEBuilder HiFi DNA Assembly Master Mix was used for the vector construction. For the generation of Tg[mnr2b-hs:EGFP-TDP-43z], EGFP was directly fused to zebrafish *tardbp* gene (EGFP-TDP-43z) and the resulting EGFP-TDP-43z was linked to the *hsp70l* promoter (650 bp) and introduced into downstream of the *mnr2b* 5’UTR in the *mnr2b*-BAC DNA (CH211-172N16, BACPAC Resources Center) via homologous recombination using a Km^r^-resistance as essentially described ^58^. Tg[mnr2b-hs:opTDP-43h] and Tg[mnr2b-hs:opTDP-43h^A315T^] lines were generated by the same procedure except that opTDP-43h and opTDP-43h ^A315T^ were used, respectively, instead of EGFP-TDP-43z. The opTDP-43h consists of the zebrafish-codon-optimized human TDP-43 that is fused directly to the zebrafish-codon-optimized mRFP1 at the N-terminus and indirectly to CRY2olig via the linker peptide TRDISIE. opTDP-43h ^A315T^ is identical to opTDP-43h except A315T mutation (GCT > ACT). All transgenic lines were created via *Tol2*-mediated transgenesis.

#### TDP-43 knockout

For the generation of *tardbp* and *tardbpl* knockout fish, target sequences for Cas9-mediated cleavage were searched by CRISPRscan ^59^. The target sequences CAAGACTTAAAAGACTACTTcgg and CAAGACTTAAAAGACTACTTcgg, where the protospacer adjacent motifs (PAMs) are indicated by lower cases, were chosen for the generation of *tardbp-n115* and *tardbpl-n94* alleles, respectively. hSpCas9 was *in vitro*-transcribed with mMESSAGE mMACHINE Kit (Thermo Fisher Scientific, AM1340) by using pCS2+hSpCas9 plasmid as a template (a gift from Masato Kinoshita, Addgene plasmid # 51815). Wild type embryos were injected with 25 pg of sgRNA and 300 pg of hSpCas9 mRNA at the one-cell stage.

#### Rescue experiment of TDP-43 knockout fish via mRNA injection

For the expression of human and zebrafish TDP-43 and its derivatives via mRNA injection, the open reading frames of zebrafish *tardbp* (TDP-43z), zebrafish-codon optimized human TDP-43 (TDP-43h), mRFP1-tagged zebrafish *tardbp* (mRFP1-TDP-43z) and opTDP-43z was cloned into pCS2+ vector *in vitro*-transcribed with mMESSAGE mMACHINE Kit. First, we injected varied amount of TDP-43z mRNA into the offspring obtained from incrosses of parental zebrafish carrying homozygous *tardbp-n115* and heterozygous *tardbpl-n94* mutation or heterozygous *tardbp-n115* and homozygous *tardbpl-n94* mutation at the one cell stage. After investigating the presence or absence of blood flow at 36-48 hpf, all fish were subjected individually to genotyping for *tardbp-n115* and *tardbpl-n94* alleles. The uninjected *tardbp-n115 tardbpl-n94* double homozygotes displayed a swollen heart that was beating, but the blood cells were completely stacked on the yolk surface and cannot reach the heart. An injection of 150 ng of TDP-43z mRNA was the most effect effective in restoring blood flow (up to 40% of the double homozygotes) of the double homozygotes with a minimum developmental abnormality due to overexpression. Throughout the assay, we scored that the blood flow was “rescued” when any blood cell flowing through the beating heart was observed. The function of TDP-43h and TDP-43 derivatives were tested by the microinjection of 150 ng mRNA each. The *tardbpl-n115* allele was identified by performing Heteroduplex Mobility Assay (HMA) against PCR product obtained with a primer pair: *tardbp-6F3* (5’-gcc aga taa taa gag gaa gat gga-3’) and *tardbpl-6R3* (5’-tga cag tac aaa gac aaa cac cac-3’). The *tardbpl-n94* allele was similarity identified by using a primer pair: *tardbpl-4F2* (5’-caa tca ctg aat gaa tgc act ttt-3’) and *tardbpl-4R2* (5’-gtt tgc tta tac taa cct gca cca-3’).

#### Blue light illumination

Short-term (< 4 hours) light stimulations of mRFP1-CRY2olig and opTDP-43z were carried out by embedding fish in the 0.8-1 % low-melting agarose (NuSieve® GTG® Agarose, Lonza) and conducting confocal scanning with the laser with 473 nm wave length using an Olympus FV1200 microscope. The average optical power of the confocal laser was approximately 44.66 *μ*W/cm^2^. For longer-term illumination, fish were raised in 6-well dish with 8 ml E3 buffer, and the dish was placed on a blue LED panel.

#### Immunohistochemistry

For the mono- and polyubiquitinated protein staining, Tg[SAGFF73A] Tg[UAS:opTDP-43z] fish at 31.5 hpf that had illuminated with a blue light were taken out from the agarose and immediately subjected to immunohistochemistry. For short, fish were fixed for 2 hours with PBS (pH 7.4) containing 4.0 % paraformaldehyde. The mouse monoclonal antibody for mono- and polyubiquitinate conjugates (1/100, FK2, Enzo) and goat anti-mouse Alexa Fluor 488 (1:1,000, Molecular Probes) were used as primary and secondary antibodies, respectively. To detect opTDP-43z, the Anti-RFP rabbit polyclonal antibody (pAb, MBL) and goat anti-rabbit Alexa Fluor 633 (1:1,000, Molecular Probes) were used as primary and secondary antibodies, respectively.

#### Microscopic analysis

For confocal microscopy, a live embryo or larva was embedded in 0.8-1 % low-melting agarose (NuSieve® GTG® Agarose, Lonza) on a Glass Base dish (IWAKI, 3010-035) and subject to confocal microscopy using an Olympus FV1200 laser confocal microscope. Images of live embryos and larvae were acquired as serial sections along the z-axis and analyzed with Olympus Fluoview Ver2.1b Viewer, Image J and Adobe Photoshop CS6. The axon length and branching frequency were measured by Imaris Filament Tracer. A neurite with more than 5 *μ*m of length was counted as branch. Morphological analyses of CaP were restricted to the spinal segment 14-17 before 50 hpf and to 13-17 during 56–72 hpf. For the quantitation of opTDP-43z in the skeletal muscle cells, averaged change of opTDP-43 intensity in the cytoplasm (D) and nucleus (E) during the illumination (N = 8 cells). *, p = 0.0097. For quantitation of opTDP-43h fluorescence intensity over EGFP-TDP-43z in the spinal motor neurons, a z-stack image of the spinal motor column was first created by Sum Slices function of ImageJ. Then, the soma area was defined by the contour of weak cytosolic fluorescence of EGFP-TDP-43z and the fluorescent intensities of opTDP-43h and EGFP-TDP-43z in the soma area were measured for individual cells using imageJ software. Statistical analyses were performed using GraphPad Prism Software.

#### RT-PCR

The total RNA was prepared from Tg[mnr2b-hs:EGFP-TDP43z], Tg[mnr2b-hs:EGFP-TDP43z] Tg[mnr2b-hs:opTDP-43h], and Tg[mnr2b-hs:EGFP-TDP43z] Tg[mnr2b-hs:opTDP-43h^A315T^] larvae at 72 hpf (17 larvae each) by homogenizing in 1 ml of Trizol Reagent (Life Technologies). Three *μ*g of the total RNA is used for cDNA synthesis using oligo dT (SuperScriptn®III First-Strand, Invitrogen). opTDP-43h and opTDP-43h^A315T^ were detected by a primer pair against the zebrafish codon-optimized mRFP1: zmRFP1-123f (5’-TCA GAC AGC TAA ACT GAA GGT CAC-3’) and zmRFP1-633r (5’-GAC GAT GGT ATA GTC TTC GTT GTG-3’). EGFP-TDP-43z was detected by a primer pair against EGFP: EGFP-f2s (5’-CAC ATG AAG CAG CAC GAC TTC T-3’) and EGFP-r5s (5’-ACG TTG TGG CTG TTG TAG TTG T-3’). zfand5b expression was detected by a primer pair: zfand5b--133f (5’-ATA GTA CAC ACC GAA ACG GAC AC-3’) and zfand5b-772r (5’-TTA TAT TCT CTG GAT TTT ATC GGC-3’)

#### Data availability

The data that support the findings in this study are available within the article and its Supplemental Information files, and from the corresponding authors upon request.

---

**Supplementary Figure 1.**
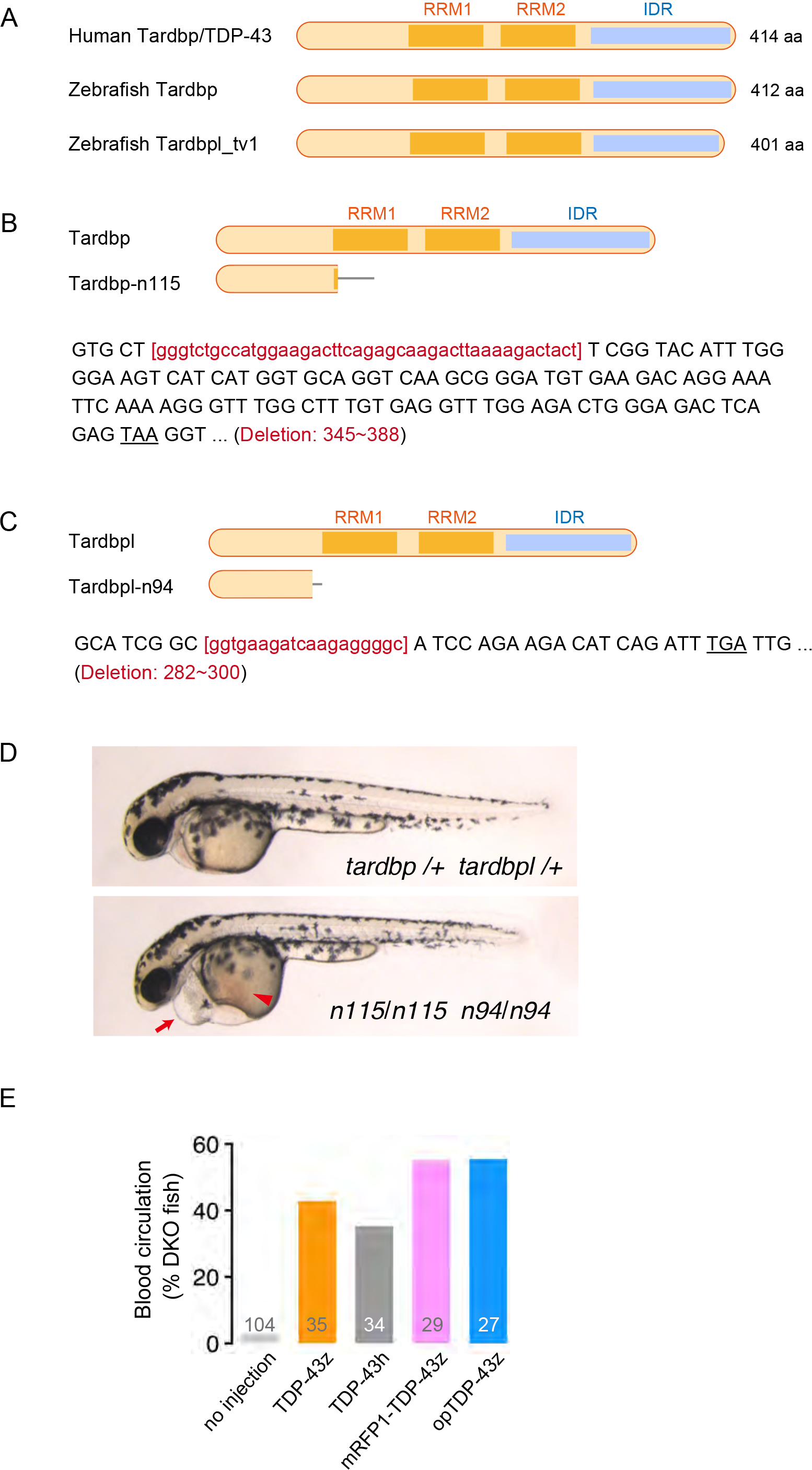
Functional validation of opTDP-43z using TDP-43 knockout fish. (A) The structure of human and zebrafish Tardbp/TDP-43 proteins. RRM, RNA-recognition motif. IDR, Intrinsically disordered region (B, C) The *tardbp-n115* and *tardbpl-n94* mutations are frame-shift deletions that cause protein truncation. The deleted nucleotides were indicated by red lower cases. The grey bars indicate ectopically added peptide due to the frame shift. (D) The lateral views of the wild type (top) and double homozygous (bottom) larvae at 48 hpf. The arrow and arrowhead indicate the swollen heart and the stacked red blood cells on the far side of the yolk, respectively. (E) Rescue rate of the blood circulation defect of the *tardbp-n115 tardbpl-n94* homozygotes (DKOs). The numbers on the histograms show the total numbers of DKOs investigated obtained from three independent microinjection experiments for each TDP-43 construct.

**Supplementary Figure 2.**
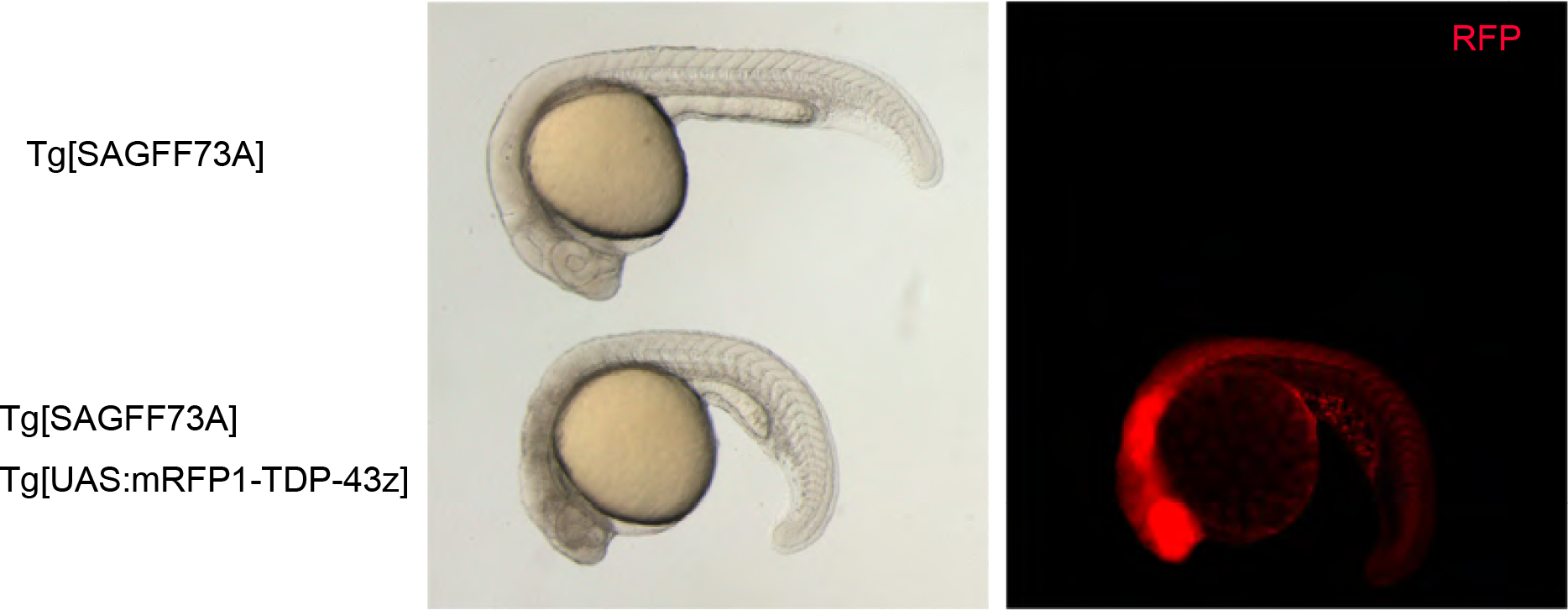
Whole-body overexpression of mRFP1-TDP-43z is toxic to zebrafish. The lateral view of Tg[SAGFF73] (top) and Tg[SAGFF73] Tg [UAS:mRFP1-TDP-43z] (bottom) embryos at 24 hpf. The RFP signal is shown in the right panel.

**Supplementary Figure 3.**
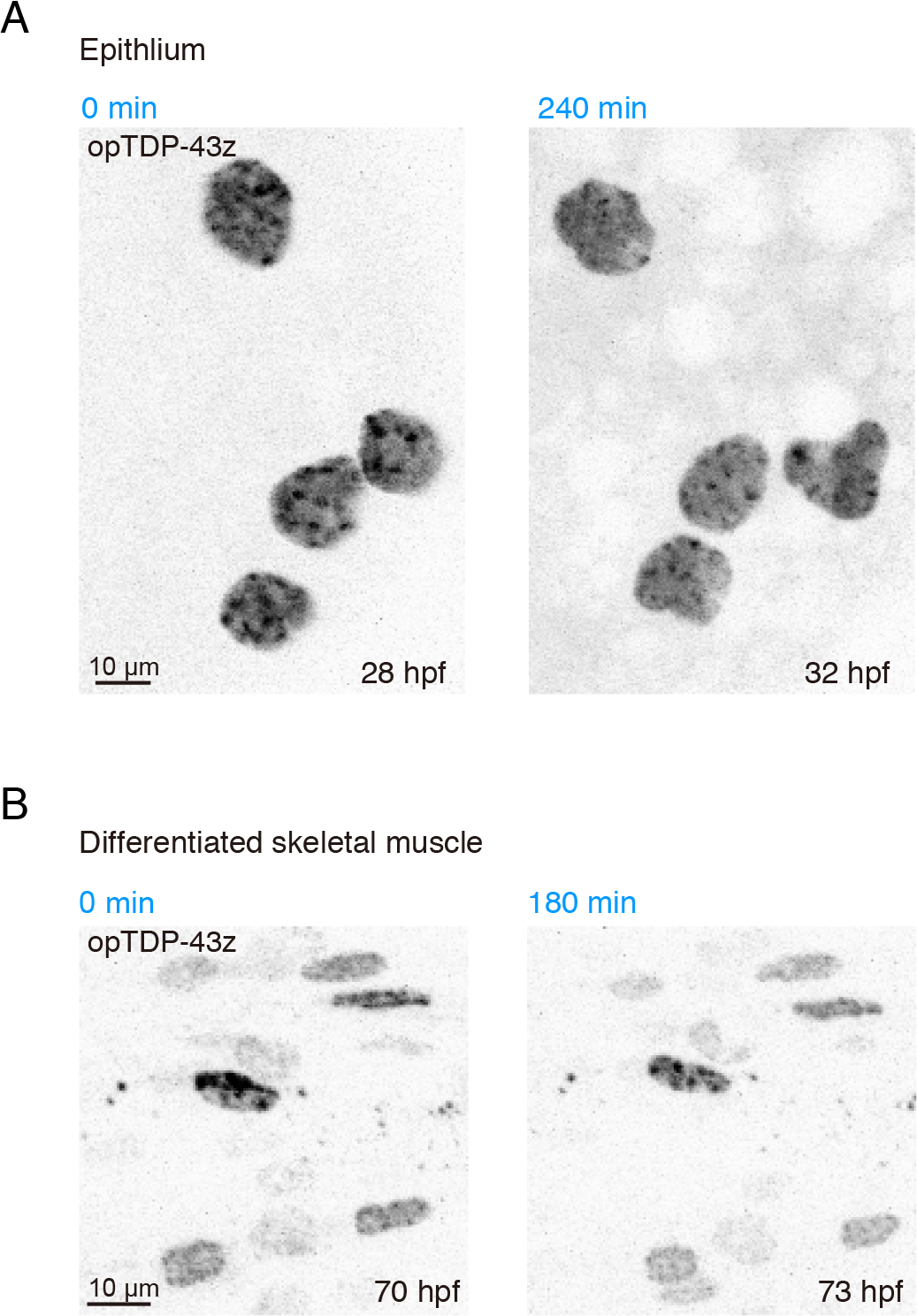
Cell-type specificity of light-dependent cytoplasmic opTDP-43z mislocalization. (A) The nuclear-enriched opTDP-43z localization was not changed by the 3-hour blue light illumination in the embryonic epithelial cells. (B) opTDP-43z localization was not changed by the 4-hour blue light illumination in the differentiated skeletal muscle cells. Cytoplasmic opTDP-43 droplets were occasionally observed independently of light illumination. These light-insensitive opTDP-43 droplets were static throughout the experiment.

**Supplementary Movie 1**

The skeletonized CaP axons of Tg[SAIG213A] Tg[UAS:EGFP] (left) and Tg[SAIG213A] Tg[UAS:EGFP] Tg[UAS:mRFP1-TDP-43z] (right). The axon branch points and terminals are indicated by red and green, respectively.

**Supplementary Movie 2**

The lateral view of CaPs of Tg[SAIG213A] Tg[UAS:mRFP1-CRY2olig] double transgenic embryo. The blue light was illuminated from 0 to 30 min (indicated by a blue dot).

**Supplementary Movie 3**

Skeletal muscle cells expressing opTDP-43z opTDP-43z illuminated with blue light in in Tg[SAGFF73A] Tg[UAS:opTDP-43z] fish at 28 hpf.

**Supplementary Movie 4**

The dorsal view of the spinal cord at the segment 14 -17 levels of a Tg[SAIG213A] Tg[mnr2b-hs:Gal4] Tg[UAS:opTDP-43z] Tg[UAS:EGFP] quadruple transgenic fish. EGFP (left) and opTDP-43z (right) were presented side by side. The transverse planes for EGFP and opTDP-43z signals are not completely matched because they are independently registered using Image J software.

**Supplementary Movie 5**

A Tg[mnr2b-hs:EGFP-TDP43z] Tg[mnr2b-hs:opTDP-43h^A315T^] (A315T) larva (focused) that had been illuminated with blue light during 2-5 dpf is lying at the bottom of the dish with normal heart beat but without the swim bladder inflation. Its sibling (Tg[mnr2b-hs:EGFP-TDP43z] larva) that experienced the same illumination procedure is capable of swimming freely with an inflated swim bladder. The movie is played in actual speed.

